# Value-driven attentional capture enhances distractor representations in early visual cortex

**DOI:** 10.1101/567354

**Authors:** Sirawaj Itthipuripat, Vy A. Vo, Thomas C. Sprague, John T. Serences

## Abstract

When a behaviorally relevant stimulus has been previously associated with reward, behavioral responses are faster and more accurate compared to equally relevant but less valuable stimuli. Conversely, task irrelevant stimuli that were previously associated with a high reward can capture attention and distract processing away from relevant stimuli (e.g. the chocolate bar in the pantry when you are looking for a nice healthy apple). While increasing the value of task-relevant stimuli systematically up-regulates neural responses in early visual cortex to facilitate information processing, it is not clear if the value of task-irrelevant distractors influences behavior via competition in early visual cortex or via competition at later stages of decision-making and response selection. Here, we measured fMRI in human visual cortex while subjects performed a value-based learning task, and applied a multivariate inverted encoding model to assess the fidelity of distractor representations in early visual cortex. We found that the fidelity of neural representations related to task-irrelevant distractors increased when the distractors were previously associated with a high reward. Moreover, this value-based modulation of distractor representations only occurred when the distractors were previously selected as targets on preceding trials. Together, these findings suggest that value-driven attentional capture begins with sensory modulations of distractor representations in early areas of visual cortex.

## Introduction

In most real-world situations, stimuli that are visually salient—such as a camera flash in a theater, or a green object in a sea of red—automatically capture attention[1–4]. Likewise, distractors that are distinguished only by their value, not their visual salience, also capture visual attention—even on occasions when high-value distractors are completely irrelevant and unactionable (e.g., a driver runs a red light because they get distracted by luxury sports car)[5–10]. In the laboratory, the value associated with an irrelevant distractor interferes with the processing of task-relevant visual information, resulting in increased response times (RTs) and sometimes reduced accuracy in a variety of tasks ranging from simple visual discrimination to more complex scenarios in which the value of multiple competing items must be compared[5–8,10–17]. Importantly, these behavioral effects of value-based attentional capture are overexpressed in patients with attention-deficit hyperactivity disorder and addiction[18,19]. While previous work has shown that the value of task-relevant visual information increases neural activity in areas of early visual cortex [20–26], it is unclear how the learned value of irrelevant distractors modulates cortical responses in these regions.

To examine this, we recruited human participants to perform a value-based decision-making task and measured their brain activity in visual cortex using functional magnetic resonance imaging (fMRI). Subjects were required to select one of two task-relevant options while ignoring a third irrelevant and unactionable distractor that was rendered in a color that had been previously associated with a variable level of reward. We hypothesized that the previously assigned value of the distractor color would modulate evoked responses in early visual cortex, and that this reward-based modulation would be specific to the spatial location of the distractor stimulus. To evaluate spatially selective modulations, we used an inverted encoding model (IEM) to reconstruct a representation of each stimulus using activation patterns of hemodynamic responses from retinotopically organized visual areas V1, V2, and V3. We found that distractors previously associated with a high value slowed choice RTs. Distractors were also represented with higher fidelity in extrastriate visual areas V2 and V3. Importantly, these value-based modulations of behavior and of neural representations depended on target selection history. That is, the effect of distractor value on behavioral and neural data only occurred when the color of the distractor matched the color of a recently selected target. Together, these results suggest that the influence of high-value distractors on attentional capture begins with an early modulation of sensory responses, and that this value-driven attentional capture occurs when participants have learned the value associated with the visual feature of the distractor.

## Results

### High-valued distractors automatically capture attention

In the present study, we used fMRI to measure activity in retinotopically organized visual areas V1, V2, and V3 while human participants (*N*=15) performed a two-alternative value-based decision-making task with changing reward associations [6] (Figure 1). On each trial, three stimuli were presented, each rendered in a different color. Two of the stimuli were presented in fixed target locations and subjects had to choose between them. The third stimulus, termed a ‘distractor’, was presented in another fixed location that subjects could never select. Participants learned that different rewards (1 or 9 cents) were associated with the colors of visual stimuli presented at the two target locations. Importantly, the distractor was not actionable and was thus completely irrelevant with respect to evaluating the relative value of the two possible targets. Across trials, the colors of the targets and the distractor changed randomly so that the distractor color on a given trial could match the color of a previously selected target that yielded either a low or a high monetary reward. Additionally, the pairings between color and reward changed across mini-blocks of 8 trials, so that values assigned to different colors could be counterbalanced. Thus, for behavioral and fMRI analyses, we sorted trials based on incentive values assigned to the colors of distractors (i.e., low- or high-valued distractor). The incentive value was always defined. However, a given color may not have been selected on previous trials. Therefore, the current value of the distractor was not always known to the participant. We thus examined the ‘selection history’ of the current distractor color by coding whether it was selected as a target in the previous 3 trials (i.e., selected or unselected; see Materials and Methods).

**Figure 1.**
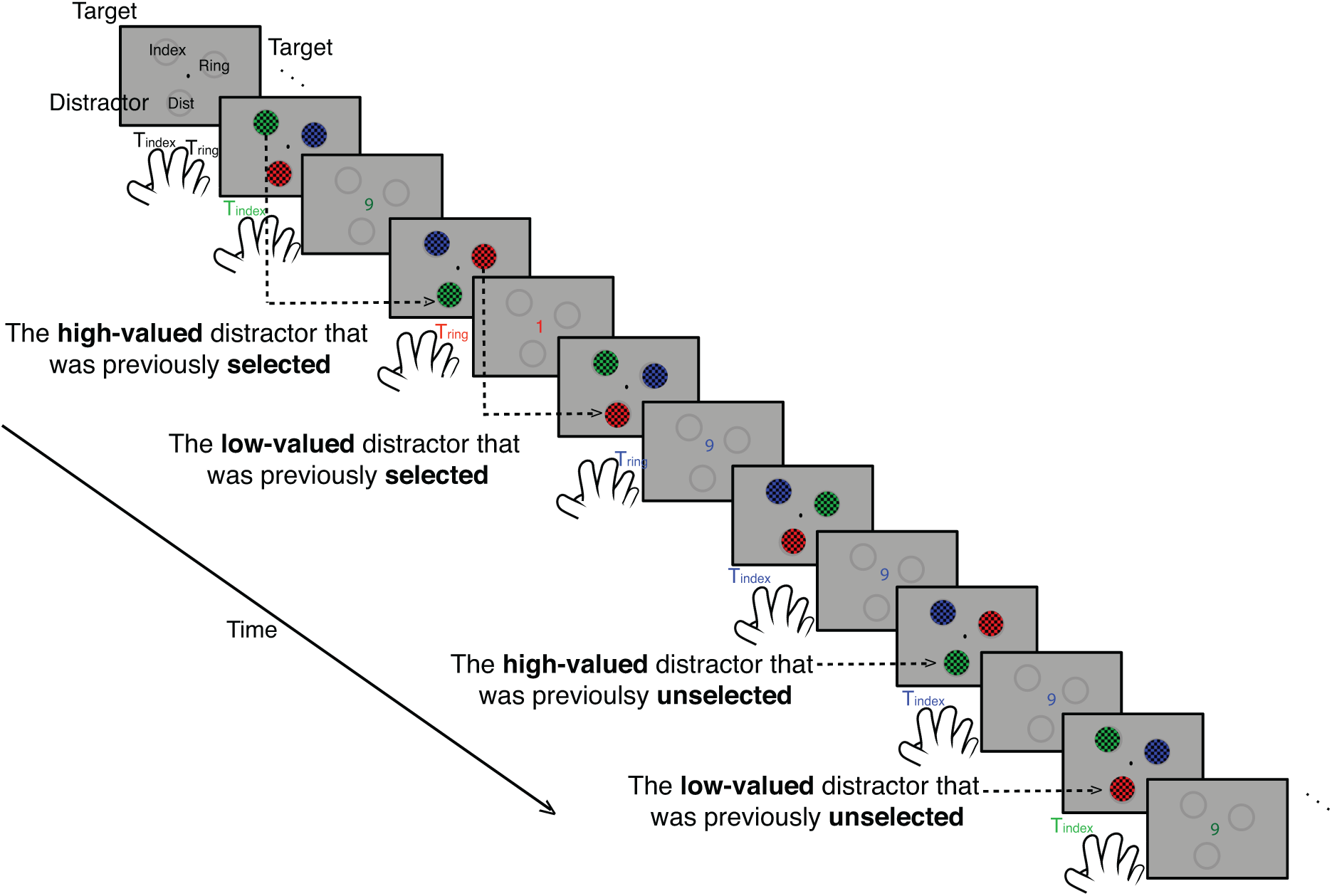
Value-based decision-making task. Participants selected one of the two target stimuli to learn values associated with their colors, while ignoring a task-irrelevant distractor that could never be selected and was thus unactionable. Across trials, the colors of the targets and the distractor changed randomly so that the distractor color on a given trial could match the color of a previously selected target that yielded either a low or a high monetary reward (i.e., low- or high-valued distractor).

Overall, subjects selected higher valued targets more often than lower valued targets (Figure 2A, p ≤ 1×10^−6^, 2-tailed, resampling test). This indicates that subjects were able to learn the values assigned to the different colors. Next, we fit the choice preference data as a function of differential target value with a cumulative Gaussian function (Figure 2B). We found no effect of distractor value (high – low distractor value) on these fit parameters on trials where the current distractors were previously selected (p’s = 0.9420 and 0.0784 for *sigma* and *mu*, respectively, 2-tailed) or unselected (Figure 2B; p’s = 0.5637and 0.8206 for *sigma* and *mu*, respectively, 2-tailed). The null distractor value effect in the choice preference data is consistent with a large body of literature demonstrating smaller and more variable distractor value effects on task accuracy [11,27,28].

**Figure 2.**
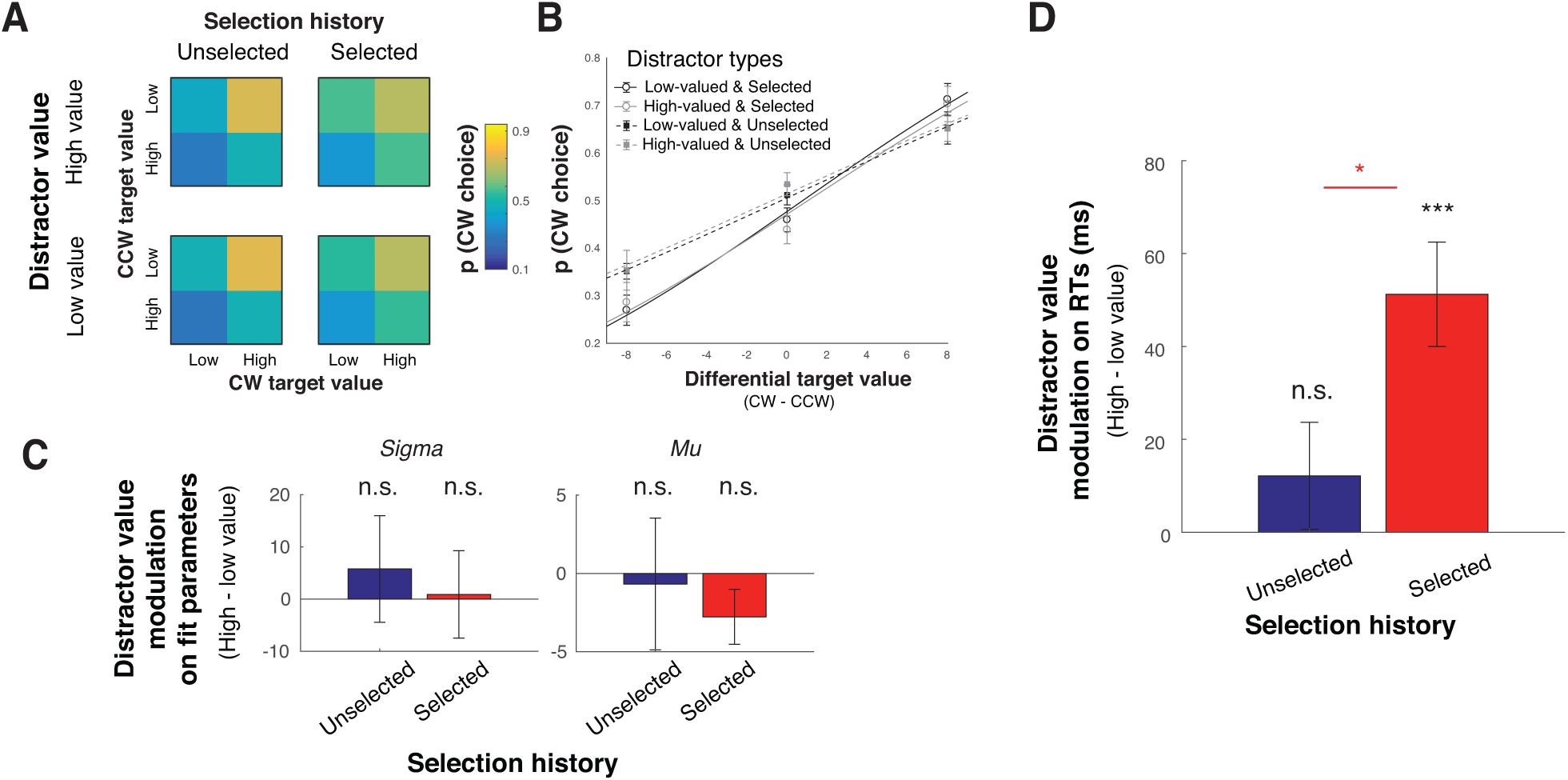
High-valued distractors increased response times. (A) Choice preference for high-valued targets for different distractor types. CW and CCW targets are targets located clockwise and counter-clockwise to the distractor location, respectively. (B) The same choice preference data, overlaid with the best fit cumulative Gaussian function (see Table 1). (C) Distractor value modulation (high – low distractor value) of regression parameters that explain choice preference functions in (B) (also see Table 1). Overall, we observed no distractor value modulation on choice preference functions: none of the regression parameters changed with distractor value in trials where the current distractor was previously selected or unselected. (D) Unlike choice preference data, we observed a robust distractor value modulation on RTs. The RT effect was significant only for trials where the distractor was previously selected. Black *** shows a significant distractor value modulation compared to zero with p < 0.001 (2-tailed; resampling test). Red * shows a significant difference between trials where the current distractors were previously selected and unselected with p < 0.05 (1-tailed). All error bars show ±1 standard error of the mean (SEM).

While there was no distractor value modulation on the choice preference data, RTs differed significantly across different distractor types (Table 1). We observed a significant effect of distractor value (high – low distractor value) on RTs on trials where the current distractor was previously selected (Figure 2D; p ≤ 1×10^−6^, 2-tailed). However, there was no distractor value modulation on trials where the current distractors were previously unselected (p = 0.2756, 2-tailed). Moreover, the magnitude of the distractor value modulation was significantly higher for the current distractor that was previously selected vs. unselected (p = 0.0102, 1-tailed). These RT results show that the distractor value captures attention, leading to a relative increase in the speed with which subjects processed task-relevant targets [5–8,13–17].

**Table 1.** Cumulative Gaussian parameters describing choice preference data and response times (RTs) for different distractor types shown in Figure 2.

| Behavioral measurements | Distractor types: Distractor value & Selection history (mean ± SEM) | | | |
| --- | --- | --- | --- | --- |
|  | Low & Unselected | High & Unselected | Low & Selected | High & Selected |
| *Sigma* | 23.99±8.01 | 29.87±9.48 | 32.66±7.09 | 33.56±7.81 |
| *Mu* | 0.85±2.41 | 0.17±2.80 | 1.31±1.58 | -1.46±1.13 |
| RTs (ms) | 600±20 | 612±15 | 592±18 | 643±19 |

### The reward history of distractors modulates neural representations in early visual cortex

To examine the influence of the distractor value on spatially specific distractor- and target-related neural representations in early visual cortex, we employed a multivariate analysis of fMRI data – an inverted encoding model (IEM; Materials and Methods; Figure 3) [20,29–31]. The IEM exploits the spatial tuning of neuronal populations in visual cortex to reconstruct representations of target and distractor stimuli based on population-level activity measured via fMRI. As expected, we found that these reconstructions peaked at the center of each of the three locations (Figure 4A; sorted as unselected target, selected target, and distractor). Qualitatively, the reconstructed activation at the distractor location was highest when the distractor colors matched the target colors that had been selected (i.e., selected distractors) and rewarded with a higher value in the previous trials (i.e., the high-valued & previously selected distractor, the top right of the Figure 4A), compared to all the other distractor conditions.

**Figure 3.**
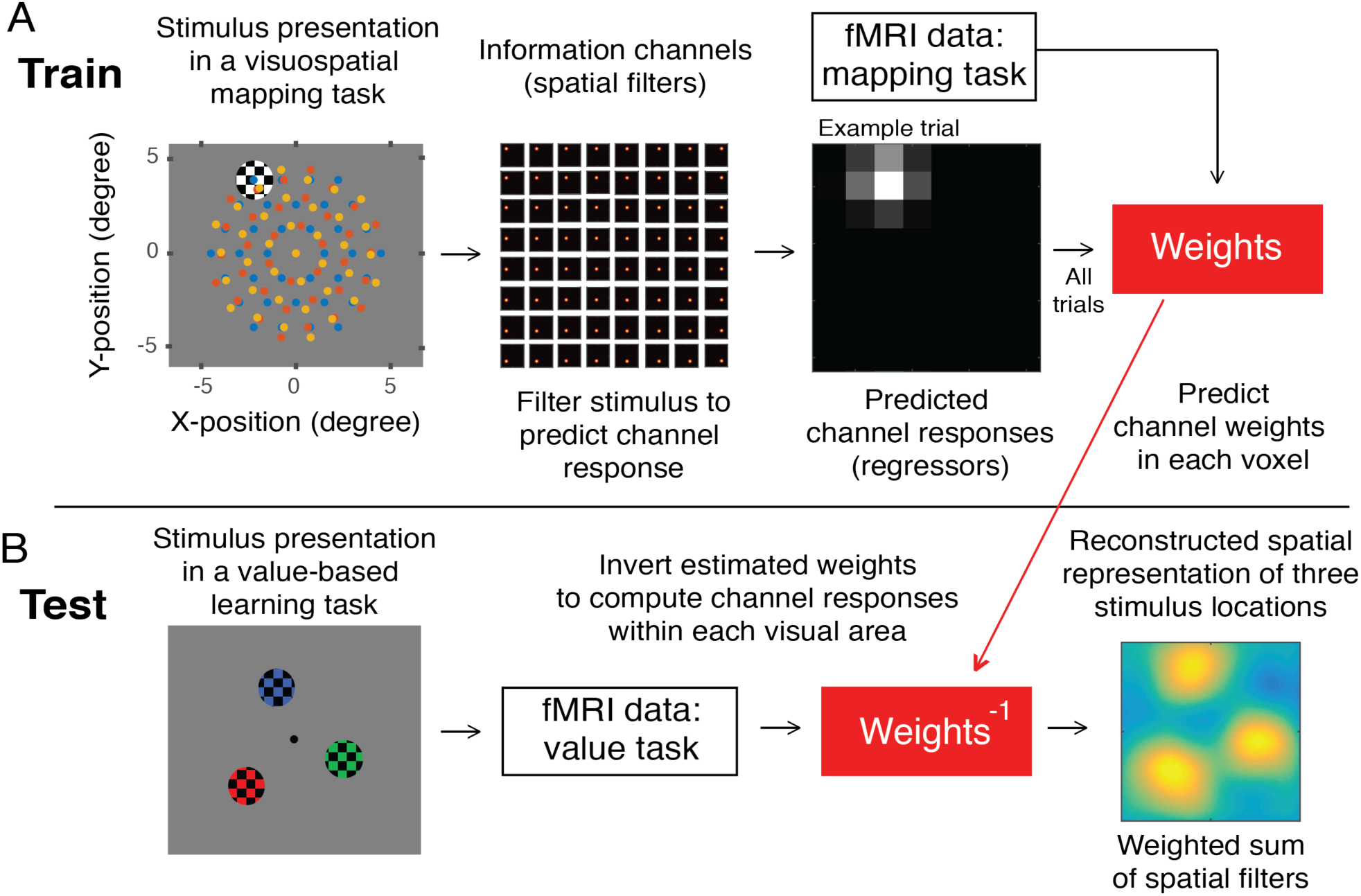
Quantifying stimulus representations with an inverted encoding model (IEM). (A) The IEM was trained using fMRI data from the visuospatial mapping task, where flickering-checkerboard mapping stimuli were randomly presented at each of 111 locations (center locations shown in blue, red, and yellow dots in the first panels; these dots were not physically presented to participants). We filtered individual stimulus locations using 64 Gaussian-like spatial filters to predict channels responses for each trial. We then use the predicted channel responses and fMRI data of all trials to predict channel weights for each voxel within each visual area. (B) The IEM was tested using fMRI data from the value-based learning task (an independent dataset). We inverted the estimated channel weights to compute channel responses within each visual area, resulting in a spatial reconstruction centered at three stimulus locations in the value-based learning task.

**Figure 4.**
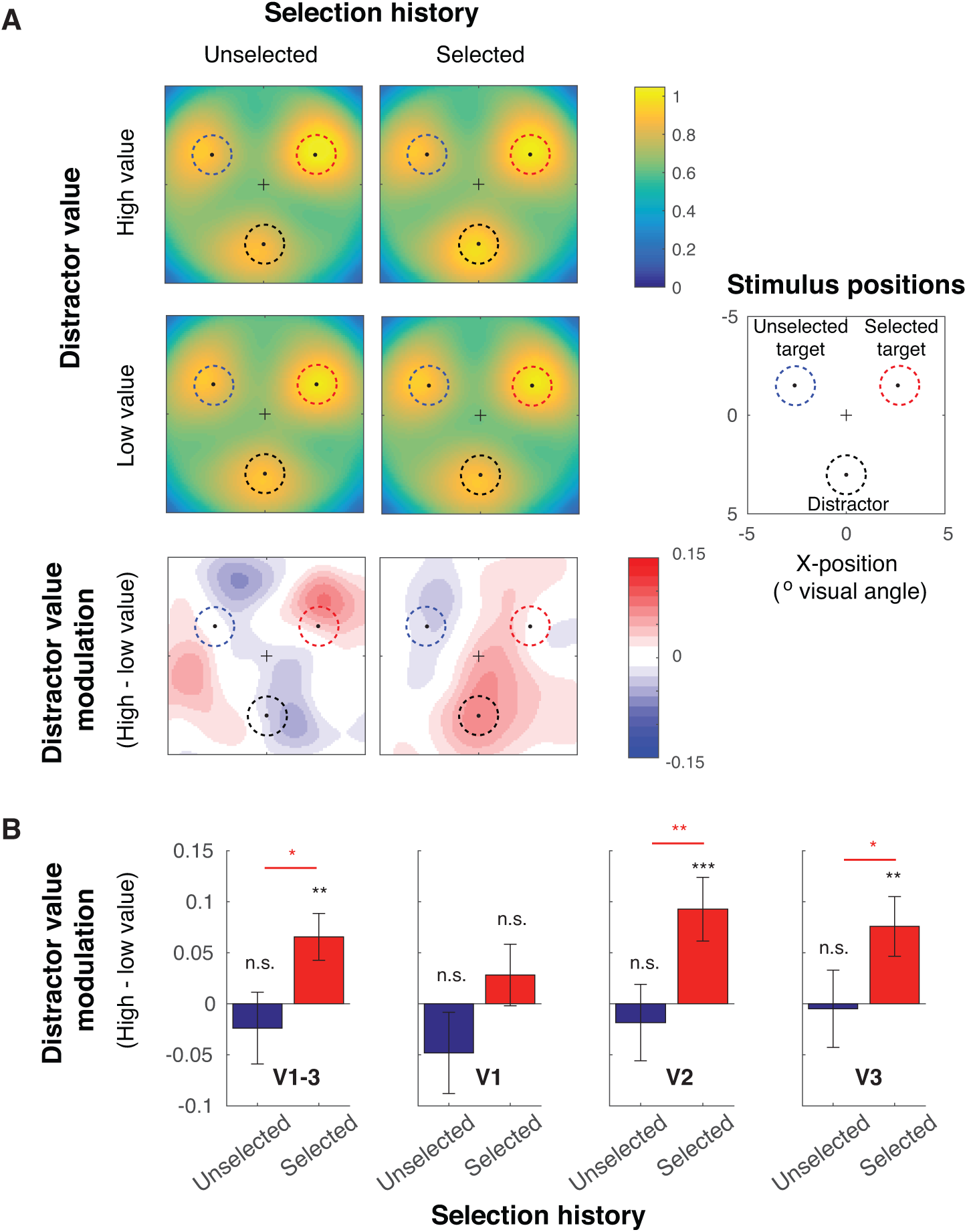
Distractor value boosted the activation of distractor representations in early visual cortex. (A) Averaged spatial reconstructions of the selected target, unselected target, and distractor based on fMRI activation patterns in early visual areas (collapsed across V1-V3). The data were sorted based on the distractor value (high and low distractor value) and the selection history (previously selected and unselected; also see Online Methods). Before averaging, reconstructions were rotated so that the positions of each respective stimulus type were in register across subjects. In each color plot, a black dot marks the location of the central fixation, and three surrounding dots at 30°, 150°, 270° polar angle indicate the centers of the selected target, unselected target, and distractor locations, respectively. The bottom panels show difference plots between high and low distractor value conditions. (B) The distractor value modulation (high – low distractor value) from the reconstruction activation (averaged across black dashed circles in A). Overall, we found significant distractor value modulations in extrastriate visual areas V2 and V3, only in trials where the current distractor was previously selected. Black ** and *** show significant distractor value modulations compared to zero with p < 0.01 and p < 0.001 (2-tailed). Red * and ** show a significant difference between trials where the current distractors were previously selected and unselected with p < 0.05 and p < 0.01 (1-tailed). The stats computed for different visual areas were corrected using the Holm-Bonferroni method. All error bars show ±1 standard error of the mean (SEM). Blue, red, and black dashed circles in A represent the spatial extents of unselected targets, selected targets, and distractors, respectively.

To quantify this effect, we computed the mean activation level in the reconstructed stimulus representations over the space occupied by the distractors (Figure 4A, see Materials and Methods; also see Sprague et al., 2018). Then, we used a non-parametric resampling method (i.e., resampling subjects with replacement) to evaluate the impact of distractor value (high vs. low distractor values) on the mean activation of the distractor representation. We did this separately for trials where the current distractor had been previously selected or unselected in preceding trials to determine if distractor value modulations depended on the selection history associated with the color of the distractor.

First, we analyzed the data averaged across V1-V3 (Figure 4B). We found a significant distractor value modulation (high > low value) for the distractor that was previously selected (p = 1 x10^−3^, 2-tailed) but a null result for the distractor that was previously unselected (p = 0.4956, 2-tailed). We directly evaluated this effect and found that selection history significantly increased distractor value modulation (p = 0.0243, 1-tailed). We then repeated these tests separately for individual visual areas. We found significant distractor value modulations for the previously selected distractor in extrastriate visual areas V2 and V3 (p= 0.0011 and p = 0.0052, passing the Holm-Bonferroni-corrected thresholds of 0.0167 and 0.025, respectively, 2-tailed) but not in the primary visual cortex V1 (p = 0.3318, 2-tailed). In V2 and V3, we confirmed that selection history had a significant effect on distractor value modulation (p = 0.0086 and p = 0.0374, respectively, 1-tailed). Similar to the data averaged across V1-V3, there was no significant distractor value modulation for the previously unselected distractors in any visual area (p = 0.2031, p = 0.6263, and p = 0.9230, for V1, V2, and V3, respectively, 2-tailed). In sum, we used an IEM to evaluate spatially-specific representations of behaviorally irrelevant stimuli with an associated reward history. We found that the value associated with irrelevant visual features is encoded in spatially-specific activation in early visual areas V2 and V3.

### Target selection and target value are encoded in early visual cortex

As shown in Figure 3A, stimulus representations are generally higher for selected targets compared to unselected targets. To quantify this effect, we computed the mean activation level in the reconstructed stimulus representations over the space occupied by the selected and unselected targets (Figure 5A). For the data collapsed across V1-V3, we observed a significant target selection modulation (selected > unselected targets: p = 0.0011 for data collapsed across distractor types; p’s = 0.0642, 0.0003, 0.0228, and 0.0022 for low-valued & unselected, high-valued & unselected, low-valued & selected, and high-valued & selected distractors, with the Holm-Bonferroni-corrected thresholds of 0.05, 0.0125, 0.025, and 0.0167, respectively, 2-tailed). These target selection modulations were significant in all visual areas (p’s = 0.0189, 4.600 x10^−4^ and p = 5.600 x10^−4^, V1, V2 and V3, respectively; Holm-Bonferroni-corrected, 2-tailed).

**Figure 5.**
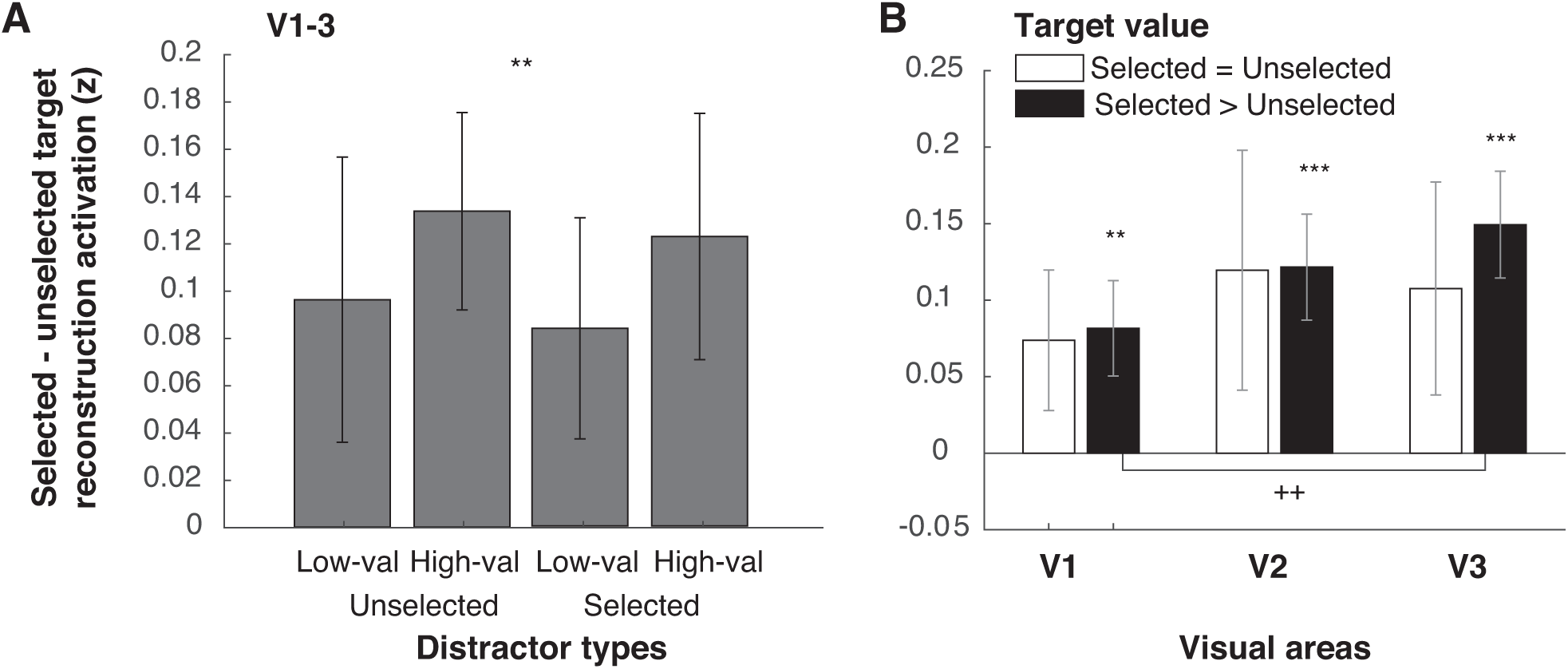
Target selection modulations in early visual areas. (A) The difference between the selected and unselected target reconstruction activation for different target types. The activation values were obtained from averaging the reconstruction activation over circular spaces spanning the spatial extents of target stimuli (red and blue dashed circles in Figure 4A). The data in (A) were collapsed across visual areas. (B) The same data as (A) but plotted separately for different target value conditions and for different visual areas. ** and *** indicate significant target selection modulations compared to zero with p’s < 0.01 and < 0.001, respectively (2-tailed). ^++^ indicate a significant difference across visual areas V1 and V3. Stats in (B) were corrected with the Holm-Bonferroni method. All sub-figures are plotted with ±1 SEM.

Next, we evaluated the impact of distractor value on the differential activity between selected and unselected targets. We found no influence of distractor value on target representations (high- vs low-valued distractors) on trials where the current distractor was previously selected (p =0.2303, 2-tailed) or on trials where the current distractor was unselected (p = 0.4463, 2-tailed). Similar null results were also observed when the data were analyzed separately in V1, V2, and V3 (p’s = 0.1639-0.8710 and 0.0744-0.9419 for the selected and unselected conditions, 2-tailed). These are consistent with the null distractor value effects on the choice preference data (Figures 2A-B).

Previous studies have reported that the relative value of targets is encoded in early visual cortex [23–25]. To test this, we analyzed the target selection modulation data both when the selected and unselected targets had the same value (i.e., selected = unselected targets), and when the selected target had a higher value compared to the unselected target (i.e., selected > unselected targets). As shown in Figure 5B, we found significant target selection modulations only when the selected targets had a higher value compared to the unselected targets in all visual areas (p’s = 0.0055, 4×10^−6^, and 1×10^−6^, passing the Holm-Bonferroni-corrected thresholds of 0.0125, 0.0100, and 0.0083 for V1, V2, and V3, respectively, 2-tailed), but no significant target modulations when selected and unselected targets had the same value (p’s = 0.0437-0.0756, which did not pass the Holm-Bonferroni-corrected threshold of 0.0167, 2-tailed). In addition, on trials where participants selected the higher-valued target, the target selection effect was significantly stronger in V3 than V1 (p = 0.0021, passing the Holm-Bonferroni-corrected of 0.0167, 2-tailed). However, there was not a significant difference between V3 and V2 (p =0.1165, 2-tailed) or between V2 and V1 (p = 0.1274, 2-tailed). Taken together with the previous section, our results show that the encoding of target value and distractor value can occur in parallel in early areas of visual cortex.

## Discussion

Visual stimuli that are not physically salient but that are paired with high reward values are known to automatically capture attention, even when those stimuli are behaviorally irrelevant and unactionable [5–9]. While a recent study reported that neural responses associated with distractors scale with reward history [32], it is unclear if these modulations were tied specifically to the location of the distractor and whether distractor response modulations led to attenuated target responses. Using a multivariate spatial reconstruction analysis of fMRI data, we show here that retinotopically organized regions in extrastriate visual areas V2 and V3 are modulated by the reward history of irrelevant visual stimuli. Importantly, the spatial reconstructions of these stimuli indicate that reward-based modulations occur precisely at the location of the distractor and that there is little associated impact on responses to simultaneously presented targets. Taken together, our results suggest that value-based modulations may begin with the early value-based modulation of sensory responses evoked by the distractor.

At the first glance, our results seem to contradict several recent studies that observed a reward-based suppression of neural representations associated with distractors in sensory cortices [33–36]. However, in many of these studies, the reward manipulation was not specifically tied to the distractor and distractor suppression was inferred based on modulations of neural responses related to the task-relevant targets [33–35]. Thus, these recent results are actually in line with the current data, in which the reconstruction activation of selected targets was higher than unselected targets and low-valued distractors. That said, another recent study reported that a high-valued distractor induced weaker neural representations in early visual cortex compared to the low-valued distractor [36]. However, they found that this was true only when the distractor was physically more salient than the target in a perceptually demanding task[36]. They reasoned that the high sensory competition between low salience targets and high salience distractors required top-down attentional suppression of the high-valued targets[36]. However, this was not the case in the current experiment, where all stimuli were suprathreshold and matched for luminance. Thus, in the context of our experimental design, we did not find evidence for distractor suppression at either the behavioral or neural level.

In the present study, we showed that an association between reward and color can induce neural modulations in early visual areas V1 – V3. This is somewhat surprising given evidence that neurons in higher visual areas, such as V4, V8, VO1, and inferior temporal cortex, are selectively tuned to chromatic information and responsible for processing color-based top-down modulations [29,37–42]. We suggest that value-based modulations in early visual areas may reflect top-down feedback signals from these higher visual areas, where the association between color and reward might be computed. Related to this idea, we found significant distractor value modulations only in extrastriate visual cortex but not in V1, which may reflect a reentrant signal backpropagated to earlier visual areas. The more robust effects in higher visual areas were also observed for the task-relevant target reconstructions, consistent with previous reports [20,30,31,43,44]. Overall, this pattern of data supports theoretical frameworks suggesting that visual cortex operates as a priority map which indexes the rank-ordered importance of different sensory inputs [20,23–25,30,31,33,34,45,46]. That said, the assumption that the color-reward association can only be computed in higher visual areas has to be considered with caution, because studies have also found that primary and extrastriate visual areas contain neuronal populations with an inhomogeneous spatial distribution of color selectivity [47,48].

In summary, we demonstrate that the learned value of irrelevant distractors automatically captures attention and that this interferes with the processing of relevant visual information. This value-based attentional capture results in increased RTs and heightened distractor representations in retinotopically organized areas of extrastriate visual cortex. Together, our findings suggest that value-driven attentional capture begins with early sensory modulations of distractor representations in visual cortex. Moreover, the modulations of both relevant targets and irrelevant distractors supports a recent re-framing of the classic dichotomy between bottom-up and top-down biasing factors in favor of a trichotomy that emphasizes a crucial role of learned reward history on the processing of relevant and irrelevant visual information [9].

## Materials and methods

### Participants

Sixteen neurologically healthy human observers with normal color vision and normal or corrected-to-normal acuity participated in the present study. Participants were recruited from the University of California, San Diego (UCSD) community and all participants provided written informed consent as required by the local Institutional Review Board at UCSD (IRB# 081318). They then completed one scanning session of the main experiment and one or two sessions of retinotopic mapping scans. Participants were compensated 20 dollars per hour in the scanner with additional monetary rewards that scaled with their behavioral performance in the value-based learning task (mean 13.13 dollars, SD 0.74). Data from one subject were excluded because of excessive movement artifacts during the retinotopy scans (>3 mm movement in more than half of the scans), leaving a total of 15 participants in the final analysis (age range 20 − 34 years old, mean age = 24.6, ±4.29 SD).

### Stimuli and tasks

Visual stimuli were rear-projected onto on a 115 cm-wide flat screen placed ∼440 cm from the participant’s eyes at the foot of the scanner bore using a LCD projector (1024×768, 60 Hz, with a grey background, luminance = 8.68 cd/m^2^). The behavioral paradigms were programmed and presented via a laptop running Windows XP using MATLAB (Mathworks Inc., Natick, MA) and the Psychophysics Toolbox [49,50].

#### Value-based decision-making task

We adopted a value-based decision-making task that we recently used to show a robust effect of distractor reward history on behavior [6]. Each block started with an instruction period, telling participants the locations of the two targets and the location of the irrelevant distractor. The position of each stimulus was indicated by different letter strings located inside three circular placeholders equally spaced from one another (120° polar angle apart with an eccentricity of 3.02° visual angle; Figure 1). The placeholders remained visible for the entire run so that participants knew the precise target and distractor locations. The instruction period was followed by experimental trials where three physically isoluminant checkerboard stimuli of different colors were presented (black paired with red, green, and blue, radius of 1.01° visual angle, and spatial frequency of 1.98 cycles per degree visual angle). The stimuli were flickered on-off at 7.5 Hz for 1 sec.

Participants were instructed to choose one of the two targets to maximize their reward, and were told that the reward value associated with each color changed across the course of the scan. The reward values associated with each stimulus color were changed every 8 trials (a mini-block). Subjects were not explicitly informed about the length of this mini-block but they were told that reward-color associations would change dynamically across a small chunk of trials. All 8 possible combinations of the three colors and two reward values (1 and 9 cents) were presented in each mini-block. The color assignments to each target and distractor stimulus were also counterbalanced within each mini-block. Trial order was pseudo-randomized so that the colors of the visual stimuli at three stimulus locations swapped in an unpredictable fashion. The assignment of different values to each color was also randomized so that changes in color-reward associations were unpredictable.

Participants were instructed to choose one of the two targets using two fingers on the right hand, as indicated in a diagram displayed before the run started (Figure 1). Importantly, the distractor could never be chosen and was thus choice-irrelevant. After a 1.25 sec delay following the offset of the stimulus array, participants received visual feedback indicating the value associated with the chosen target color (‘1’ or ‘9’; feedback duration = 0.25 sec). If a response was not given before the stimulus offset, they would receive a letter ‘M’ (“miss”) to indicate that no reward was earned on that trial. On a random 20% of trials, rewards were withheld to encourage participants to explore and learn the value of each color (done independently for each of the two targets). ‘0’ cents were given in these trials indicating that participants received no reward. The feedback period was followed by a blank inter-trial interval with a central fixation for 1.5 sec.

Participants completed 6 total blocks with the distractor location remaining stable for 2 consecutive blocks to ensure that participants knew the exact position of the distractor stimulus. Across all blocks the distractor location was counterbalanced across the 3 possible stimulus positions. Each block lasted 4 min 57 sec and contained 48 experimental trials and 20 pseudorandomly interleaved null trials. There was a blank period of 9 sec at the end of each block. We counterbalanced stimulus configurations across participants to ensure our results were not influenced by any spatial bias. To sample data from the entire circular space across subjects, the stimulus arrays were rotated by 30° polar angle to form four configurations (15°-135°-255°, 45°-165°-285°, and 75°-195°-315°, and 105°-225°-345°) and these four configurations were counterbalanced across subjects. Each subject viewed 1 of these 4 configurations for their entire scanning session.

#### Visuospatial mapping task

Participants also completed 4-7 blocks of a visuospatial mapping task (one completed 4 blocks, one completed 7 blocks, and the rest completed 6 blocks). The data from this task were then used as an independent data set to train an inverted encoding model (IEM) that was used to reconstruct spatial representations of the targets and distractors in the value-based learning task (see the analysis section below for more details). Participants were instructed to fixate centrally and to covertly attend to a checkerboard stimulus rendered at 100% Michelson contrast that pseudo-randomly appeared at different locations on the screen (3 sec duration; the same size, spatial frequency, and flicker frequency as the stimulus in the value-based learning task). The participant’s task was to detect a rare and brief dimming in contrast (19.57% target trials; 0.5 sec duration; occurring between 0.5-2 seconds after stimulus onset). On each trial, the checkerboard stimulus was presented at one of 37 locations on a triangular grid (1.50° visual angle between vertices), covering a visual space that overlapped with the stimulus locations in the value-based learning task (the first panel in Figure 3A). To smoothly cover the entire circular space, we randomly rotated the entire triangular grid around its center by 0°, 20°, or, 40° polar angle across different runs (blue, yellow, and red dots in the first panel in Figure 3A), so there were 111 different stimulus locations in total (see similar methods in Sprague et al., 2018). On each run, there were a total of 37 non-targets (1 repeat per location) and 9 targets. Target locations were pseudo-randomly drawn from the 37 locations (never repeated within each block). The magnitude of the contrast change was adjusted across trials so that accuracy was at ∼76% (mean hit = 77.95%, SD = 12.23%). Each stimulus presentation was followed by an ITI of 2-5 sec (uniformly distributed). We pseudo-randomly interleaved 10 null trials and included a blank period of 8.2 sec at the end of the block. Each block lasted 6.28 minutes.

### Behavioral analysis

We first sorted trials from the main value-based decision-making task based on target selection (i.e., target type: selected and unselected), target value (low and high value), distractor value based on previous target rewards associated with the color of the distractor (low and high value), and selection history (i.e., whether the distractor was previously unselected or selected at least once in 3 preceding trials). We chose the 3-back analysis window because it yielded the most balanced number of trials between individual conditions. That said, an analysis using a window covering 1 or 2 previous trials yielded qualitatively consistent results. Note that because of the boundary between miniblocks (every 8 trials where value-color assignments were the same), we could only go back 1 and 2 trials for the 2^nd^ and 3^rd^ trials, respectively. We excluded data from the 1^st^ trial of every 8 trials in each mini-block to reduce the spill-over effect from different sets of value-color assignments.

Next, we examined subjects’ choice preference. To do so, we labeled targets located clockwise (CW) and counter-clockwise (CCW) to the distractor CW and CCW targets and computed the probability that participants chose CW over CCW targets and plotted as a function of CW target value and CCW target value (Figure 2A). Next, we plotted the choices as a function of differential target value (CW – CCW) separately for different distractor values and fit individual subjects’ data with the cumulative Gaussian function (Figure 2B). Specifically, we estimated the mean (or *mu*) and the standard deviation (or *sigma*) of the cumulative Gaussian function that best fit the choice preference data derived from different distractor values (see Table 1 for mean and SEM)[6]. To test distractor value modulations on these parameters, we computed the bootstrap distribution of the difference in these parameters between the high and low distractor value conditions (i.e., resampling subjects with replacement for 100,000 iterations) and calculated the percentage of values in this distribution that were larger or smaller than zero to yield a 2-tailed p-value. We performed this statistical analysis separately for previously selected and unselected distractors (see above).

Finally, we examined the effect of distractor value on RTs. First, we computed the mean RTs across different distractor values for individual subjects. Then, we computed the bootstrap distribution of the RT difference between the high and low distractor value conditions (i.e., resampling subjects with replacement for 100,000 iterations) and calculated the percentage of values in this distribution that were larger or smaller than zero (a 2-tailed p-value). We performed this statistical analysis separately for previously selected and unselected distractors. We then compared whether the effect of distractor value was significantly larger in the selected condition than the unselected condition by a similar procedure that compared the two bootstrap distributions. Since we only observed significantly larger RT differences for previously selected targets, we knew the expected direction of the effect and therefore computed a 1-tailed p-value.

### fMRI analysis

#### fMRI acquisition

All MRI data were acquired on a GE 3T MR750 scanner at the Keck Center for Functional Magnetic Resonance Imaging (CFMRI) at UCSD. Unless otherwise specified, all data were collected using a 32-channel head coil (Nova Medical). We acquired functional data using a multiband echo-planar imaging (EPI) protocol (Stanford Simultaneous Multi-Slice sequence). We acquired 9 axial slices per band at a multiband factor of 8, for 72 total slices (2×2×2 mm^3^ voxel size; 800 ms TR; 35 ms TE; 35° flip angle; 104×104 cm matrix size). Prior to each functional scan, 16 TRs were acquired as reference images for image reconstruction. Raw k-space data were reconstructed into NIFTI format image files on internal servers using scripts provided by CFMRI. In each session, we also acquired forward and reverse phase encoding blips to estimate the susceptibility off-resonance field [51]. This was used to correct EPI signal distortion using FSL topup [52,53], the results of which was submitted to further preprocessing stages described below. In each session, we also acquired an accelerated anatomical using parallel imaging (GE ASSET on a FSPGR T1-weighted sequence; 1×1×1 mm^3^ voxel size; 8136 ms TR; 3172 ms TE; 8° flip angle; 172 slices; 1 mm slice gap; 256×192 cm matrix size). This same-session anatomical was coregistered to the functional data. It was also coregistered to a high-resolution anatomical from the retinotopic mapping session(s).

#### Retinotopic mapping

To identify regions of interest (ROIs) in early visual cortex, we used a combination of retinotopic mapping methods. Individual participants completed meridian mapping (1-2 ∼5-min blocks), where they saw flickering checkerboards “bowties” along the horizontal and vertical meridians while fixating centrally. They also completed several scans of a polar angle mapping task (4-6 ∼6-min blocks) where participants covertly attended to a rotating a checkerboard wedge and detected brief contrast changes (see details in Sprague and Serences, 2013; Vo et al., 2017). We identified retinotopically organized regions of visual areas V1, V2, and V3 using a combination of retinotopic maps of visual field meridians and polar angle preferences for each voxel in these visual areas and concatenated left and right hemispheres as well as dorsal and ventral aspects of individual areas [54,55]. Visual area borders were drawn on an inflated cortical surface created from a high-resolution anatomical scan (FSPGR T1-weighted sequence; 1×1×1 mm^3^; 8136 ms TR; 3172 ms TE; 8° flip angle; 172 slices; 1 mm slice gap; 256×192 cm matrix size) collected with an 8-channel head coil.

#### fMRI data preprocessing

Analysis was performed in BrainVoyager 20.2 (Brain Innovation, Maastricht, The Netherlands) supplemented with custom analysis scripts written in MATLAB R2016a (The Mathworks Inc., Natick, Mass). Using the distortion-corrected images, we first performed slice-time correction, affine motion correction, and temporal high-pass filtering. Then the functional data were coregistered to the same-session anatomical and transformed to Talairach space. Each voxel’s timecourse was *z*-scored within each run. We then built a design matrix with individual trial predictors convolved with a double-gamma HRF (peak = 5 s, undershoot peak = 15 s; response undershoot ratio = 6; response dispersion = 1; undershoot dispersion = 1). We also included a baseline predictor. This allowed us to calculate single-trial beta weights using a general linear model (GLM). These beta weights served as input to the IEM described below.

#### Inverted encoding model (IEM)

In order to create the reconstructions of target and distractor stimuli in the value-based learning task from individual ROIs, we employed an IEM for retinotopic space (see Figure 3; also see Brouwer & Heeger, 2009; Sprague et al., 2018; Sprague & Serences, 2013; Vo, Sprague, & Serences, 2017). First, we computed a spatial sensitivity profile (i.e., an encoding model) for each voxel, parameterized as a weighted sum of experimenter-defined information channels (i.e. spatial filters in second panel of Figure 3A) using an independent training data set acquired from the visuospatial mapping task (using only non-target trials). Then, we inverted the encoding models across all voxels to compute weights on the spatial information channels and used these weights to transform the fMRI data from the value-based learning task into an activation score. Specifically, the activation of each voxel is a weighted sum of 64 Gaussian-like spatial information channels arrayed in an 8 × 8 rectangular grid (see the second panel of Figure 3). The filter centers were equally spaced by 1.43° visual angle with full-width half-maximum of 2° visual angle). The Gaussian-like function of each filter is described by:

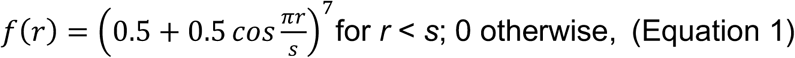

where *r* is the distance from the filter center and *s* is a size parameter indicating the distance between filter centers at which the filter returns to 0. We set values greater than *s* to 0 (*s* = 5.0332), resulting in a smooth filter at each position along the grid [30].

We then define the idealized response of the information channels for each given training trial. To do this, we multiplied a discretized version of the stimulus (*n* trials x *p* pixels) by the 64 channels defined by Equation 1 (*p* pixels x *k* channels). We then normalized this result so that the maximum channel response is 1. This is *C_1_* in the following equation:

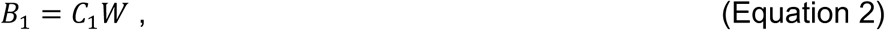

where *B_1_* (*n* trials × *m* voxels) is the measured fMRI activity of each voxel during the visuospatial mapping task (i.e., beta weights, see fMRI Preprocessing section), *C_1_* (*n* trials × *k* channels) is the predicted response of each spatial filter (i.e., information channel normalized from 0 to 1), and *W* is a weight matrix (*k* channels × *m* voxels) that quantifies the contribution of each information channel to each voxel. Next, we used ordinary least-squares linear regression to solve for *W* with the following equation:

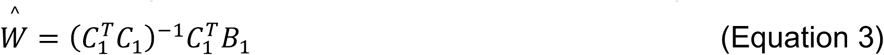

Here, 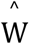 represents all estimated voxel sensitivity profiles, which we computed separately for each ROI. Next, we used 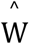 and the measured fMRI activity of each voxel (i.e., beta weights) during each trial of the value-based learning task to estimate the activation of each information channel using the following equation (see Figure 3B):

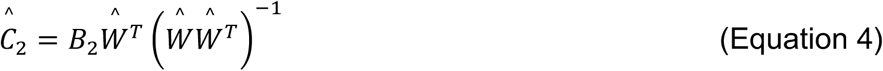

Here, 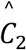 represents the estimated activation of each information channel (*n_2_* trials × *k* channels), which gives rise to the observed activation pattern across all voxels within that ROI (*B_2_*, *n_2_* trials × *m* voxels). To visualize and co-register trials across three stimulus locations, we computed spatial reconstructions by multiplying the spatial profile of each filter by the estimated activation level of the corresponding channel (i.e. computing a weighted sum; the last panel of Figure 3B). We rotated the center position of the spatial filters on each trial of individual participants such that the resulting 2D reconstructions of the target and distractor stimuli share common positions across trials and participants (CCW target, CW target, and distractor locations centered at 30°, 150°, and 270° polar angle, respectively; 3.02° visual angle from the center of the 2D reconstruction). Next, we sorted trials based on choice selection (selected and unselected) and target value (1 and 9 cents) and the reward history of the distractor (zero, low, and high) in the same way as we did for the behavioral analysis. Then we flipped all spatial reconstructions left to right on trials where the selected target location was on the left (150°) so that the unselected and selected targets always shared common locations on the left and right of the reconstruction, respectively (150° and 30°). This step did not change the position of the distractor, so it stayed at 270° polar angle. Finally, we averaged the 2D reconstructions across trials with the same trial types for individual participants and then averaged those reconstructions across participants, resulting in the grand-average spatial reconstructions shown in Figure 4A.

#### fMRI statistical analysis

Following a previous approach [20,56], we extracted the reconstruction activation for each trial type in individual participants by averaging the data within the circular space spanning the entire area of individual stimuli. This was used as our “reconstruction activation” measure. Like the behavioral analyses, all statistical analyses were conducted by resampling relevant values from each subject with replacement for 100,000 iterations and comparing these values across resampling iterations

First, we examined the distractor value modulation on the distractor reconstruction activation for data averaged across V1-V3. To do so, we computed the bootstrap distribution of the difference of the distractor reconstruction activation between the high and low distractor value conditions and calculated the percentage of values in this distribution that were larger or smaller than zero (2-tailed). We performed this statistical analysis separately for trials where the current distractor was previously selected and unselected in preceding trials to examine if the distractor value modulation depended on selection history. We then compared whether the effect of distractor value was significantly larger in the selected condition than the unselected condition by a similar procedure that compared the two bootstrap distributions (1-tailed to the known direction of the difference). We repeated the same statistical procedures for individual visual areas, and corrected for multiple comparisons using the Holm-Bonferroni method[57].

Next, we tested the target selection modulation on the target reconstruction activation for data averaged across V1-V3. To do so, we computed the bootstrap distribution of the difference between the selected and unselected target reconstruction activation and calculated the percentage of values in this distribution that were larger or smaller than zero (2-tailed). We first performed this on the data collapsed across all distractor types. Then we assessed the target selection modulations separately for individual distractor values and corrected for multiple comparisons using the Holm-Bonferroni method. Then, we tested for the distractor value modulation on the target selection modulation by computing the bootstrap distribution of the difference of the target selection modulations between the high and low distractor value conditions and computing the percentage of values in this distribution that were larger or smaller than zero (2-tailed). This was done separately for trials where the current distractor was previously unselected and selected in preceding trials. We repeated the same statistical procedures for individual visual areas, and corrected for multiple comparisons using the Holm-Bonferroni method.

Finally, we tested whether target selection modulations depended on the relative value difference between selected and unselected targets, as suggested by previous studies[23–25]. For each target value condition (same vs different target values) and each visual area, we computed the bootstrap distribution of the difference between the selected and unselected target reconstruction activation and calculated the percentage of values in this distribution that were larger or smaller than zero (2-tailed). Here, we also corrected for multiple comparisons across different target value conditions and different visual areas using the Holm-Bonferroni method (6 comparisons). Since we found more robust target selection modulations in higher visual areas in trials where the selected and unselected targets had different values, we further tested if the target selection modulation in V3 was higher than that in V1, if the target modulation in V2 was higher than that in V1, and if the target modulation in V2 was higher than that V1. To do so, we compared the target selection modulation distributions across these visual areas (1 tailed, due to the known direction of the difference), and corrected for multiple comparisons using the Holm-Bonferroni method.

## Acknowledgements

Funding provided by NEI R01-EY025872 to J.T.S., a James S. McDonnell Foundation Scholar Award to J.T.S., the Howard Hughes Medical Institute International student fellowship to S.I., a Royal Thai Scholarship from the Ministry of Science and Technology Thailand to S.I., NSF GRFP to V.A.V., and NEI F32-EY028438 to T.C.S. We thank Margaret Henderson for help with data processing, Chaipat Chunharas for assistance with data collection, and Edward Vul for useful discussions.

## Author Contributions

SI conceived and implemented the experiments, collected and analyzed the data, wrote the first draft of the manuscript, and edited the manuscript. VAV collected and analyzed the data and co-wrote the manuscript. TCS conceived the experiments and co-wrote the manuscript. JTS conceived the experiments, supervised the project, and co-wrote the manuscript.

## References

1. Yantis S, Jonides J. Abrupt visual onsets and selective attention: evidence from visual search. J Exp Psychol Hum Percept Perform. 1984;10: 601–621. doi:10.1037/0096-1523.10.5.601

2. Theeuwes J, Krueger L, Chun M, Pashler H. Perceptual selectivity for color and form. Percept Psychophys. 1992;51: 599–606.

3. Egeth HE, Yantis S. Visual attention: control, representation, and time course. Annu Rev Psychol. 1997;48: 269–297. doi:10.1146/annurev.psych.48.1.269

4. Wolfe JM. Visual Search. Attention, Perception, Psychophys. 1998;20: 13–73. doi:10.1016/j.tics.2010.12.001

5. Anderson BA, Laurent PA, Yantis S. Value-driven attentional capture. Proc Natl Acad Sci U S A. 2011;108: 10367–71. doi:10.1073/pnas.1104047108

6. Itthipuripat S, Cha K, Rangsipat N, Serences JT. Value-based attentional capture influences context-dependent decision-making. J Neurophysiol. 2015;114: 560–569. doi:10.1152/jn.00343.2015

7. Hickey C, Chelazzi L, Theeuwes J. Reward changes salience in human vision via the anterior cingulate. J Neurosci. Society for Neuroscience; 2010;30: 11096–103. doi:10.1523/JNEUROSCI.1026-10.2010

8. Anderson BA. Going for it: The economics of automaticity in perception and action. Curr Dir Psychol Sci. 2017;26: 140–145. doi:10.1177/0963721416686181

9. Awh E, Belopolsky A V., Theeuwes J. Top-down versus bottom-up attentional control: A failed theoretical dichotomy. Trends Cogn Sci. Elsevier Ltd; 2012;16: 437–443. doi:10.1016/j.tics.2012.06.010

10. Libera C Della, Chelazzi L. Learning to attend and to ignore is a matter of gains and losses. Psychol Sci. 2009;20: 778–784. doi:10.1111/j.1467-9280.2009.02360.x

11. Gluth S, Spektor MS, Rieskamp J. Value-based attentional capture affects multi-alternative decision making. Elife. 2018;7: 1–36. doi:10.7554/eLife.39659

12. Moher J, Anderson BA, Song JH. Dissociable Effects of Salience on Attention and Goal-Directed Action. Curr Biol. Elsevier Ltd; 2015;25: 2040–2046. doi:10.1016/j.cub.2015.06.029

13. Hickey C, van Zoest W. Reward-associated stimuli capture the eyes in spite of strategic attentional set. Vision Res. Elsevier Ltd; 2013;92: 67–74. doi:10.1016/j.visres.2013.09.008

14. Maclean MH, Diaz GK, Giesbrecht B. Irrelevant learned reward associations disrupt voluntary spatial attention. Atten Percept Psychophys. Attention, Perception, & Psychophysics; 2016;78: 2241–2252. doi:10.3758/s13414-016-1103-x

15. Maclean MH, Giesbrecht B. Neural evidence reveals the rapid effects of reward history on selective attention. Brain Res. Elsevier; 2015;1606: 86–94. doi:10.1016/j.brainres.2015.02.016

16. MacLean MH, Giesbrecht B. Irrelevant reward and selection histories have different influences on task-relevant attentional selection. Attention, Perception, Psychophys. 2015;77: 1515–1528. doi:10.3758/s13414-015-0851-3

17. Krebs RM, Boehler CN, Egner T, Woldorff MG. The neural underpinnings of how reward associations can both guide and misguide attention. J Neurosci. 2011;31: 9752–9759. doi:10.1523/JNEUROSCI.0732-11.2011

18. Sali AW, Anderson BA, Yantis S, Mostofsky SH, Rosch KS. Reduced value-driven attentional capture among children with ADHD compared to typically developing controls. J Abnorm Child Psychol. Journal of Abnormal Child Psychology; 2018;46: 1187–1200. doi:10.1007/s10802-017-0345-y

19. Anderson BA, Faulkner ML, Rilee JJ, Yantis S, Ph D, Marvel CL, et al. Attentional bias for non-drug reward is magnified in addiction. 2014;21: 499–506. doi:10.1037/a0034575.Attentional

20. Sprague TC, Itthipuripat S, Vo VA, Serences JT. Dissociable signatures of visual salience and behavioral relevance across attentional priority maps in human cortex. J Neurophysiol. 2018;119: 2153–2165. doi:10.1152/jn.00059

21. Zhang X, Zhaoping L, Zhou T, Fang F. Neural activities in V1 create a bottom-up saliency map. Neuron. Elsevier Inc.; 2012;73: 183–92. doi:10.1016/j.neuron.2011.10.035

22. Chen C, Zhang X, Wang Y, Zhou T, Fang F. Neural activities in V1 create the bottom-up saliency map of natural scenes. Exp Brain Res. Springer Berlin Heidelberg; 2016;234: 1769–1780. doi:10.1007/s00221-016-4583-y

23. Serences JT. Value-Based Modulations in Human Visual Cortex. Neuron. Elsevier Ltd; 2008;60: 1169–1181. doi:10.1016/j.neuron.2008.10.051

24. Serences JT, Saproo S. Population Response Profiles in Early Visual Cortex Are Biased in Favor of More Valuable Stimuli. J Neurophysiol. 2010;104: 76–87. doi:10.1152/jn.01090.2009

25. Stănişor L, van der Togt C, Pennartz CMA, Roelfsema PR. A unified selection signal for attention and reward in primary visual cortex. Proc Natl Acad Sci U S A. National Academy of Sciences; 2013;110: 9136–41. doi:10.1073/pnas.1300117110

26. Baruni JK, Lau B, Salzman CD. Reward expectation differentially modulates attentional behavior and activity in visual area V4. Nat Neurosci. 2015;18: 1656–1663. doi:10.1038/nn.4141

27. Louie K, Khaw MW, Glimcher PW. Normalization is a general neural mechanism for context-dependent decision making. Proc Natl Acad Sci. 2013;110: 6139–6144. doi:10.1073/pnas.1217854110

28. Chau BKH, Kolling N, Hunt LT, Walton ME, Rushworth MFS. A neural mechanism underlying failure of optimal choice with multiple alternatives. Nat Neurosci. Nature Publishing Group; 2014;17: 463–470. doi:10.1038/nn.3649

29. Brouwer GJ, Heeger DJ. Decoding and reconstructing color from responses in human visual cortex. J Neurosci. 2009;29: 13992–14003. doi:10.1523/JNEUROSCI.3577-09.2009

30. Sprague TC, Serences JT. Attention modulates spatial priority maps in the human occipital, parietal and frontal cortices. Nat Neurosci. 2013;16: 1879–87. doi:10.1038/nn.3574

31. Vo VA, Sprague TC, Serences JT. Spatial Tuning Shifts Increase the Discriminability and Fidelity of Population Codes in Visual Cortex. J Neurosci. 2017;37: 3386–3401. doi:10.1523/JNEUROSCI.3484-16.2017

32. Anderson BA, Laurent PA, Yantis S. Value-driven attentional priority signals in human basal ganglia and visual cortex. Brain Res. Elsevier; 2014;1587: 88–96. doi:10.1016/J.BRAINRES.2014.08.062

33. Hickey C, Peelen M V. Neural mechanisms of incentive salience in naturalistic human vision. Neuron. Cell Press; 2015;85: 512–518. doi:10.1016/J.NEURON.2014.12.049

34. Hickey C, Peelen M V. Reward selectively modulates the lingering neural representation of recently attended objects in natural scenes. J Neurosci. Society for Neuroscience; 2017;37: 7297–7304. doi:10.1523/JNEUROSCI.0684-17.2017

35. Barbaro L, Peelen M V., Hickey C. Valence, not utility, underlies reward-driven prioritization in human vision. J Neurosci. 2017;37: 1128–17. doi:10.1523/JNEUROSCI.1128-17.2017

36. Gong M, Jia K, Li S. Perceptual Competition Promotes Suppression of Reward Salience in Behavioral Selection and Neural Representation. J Neurosci. 2017;37: 6242–6252. doi:10.1523/JNEUROSCI.0217-17.2017

37. Zeki SM. Functional organization of a visual area in the posterior bank of the superior temporal sulcus of the rhesus monkey. J Physiol. 1974;236: 549–573. doi:10.1113/jphysiol.1974.sp010452

38. Conway BR, Moeller S, Tsao DY. Specialized Color Modules in Macaque Extrastriate Cortex. Neuron. 2007;56: 560–573. doi:10.1016/j.neuron.2007.10.008

39. Zeki S, Bartels A. The architecture of the colour centre in the human visual brain: new results and a review *. Eur J Neurosci. 2000;12: 172–193. doi:10.1046/j.1460-9568.2000.00905.x

40. Brewer AA, Liu J, Wade AR, Wandell BA. Visual field maps and stimulus selectivity in human ventral occipital cortex. Nat Neurosci. 2005;8: 1102–1109. doi:10.1038/nn1507

41. Brouwer GJ, Heeger DJ. Categorical Clustering of the Neural Representation of Color. J Neurosci. 2013;33: 15454–15465. doi:10.1523/JNEUROSCI.2472-13.2013

42. Hadjikhani N, Liu AK, Dale AM, Cavanagh P, Tootell RBH. Retinotopy and color sensitivity in human visual cortical area V8. Nat Neurosci. 1998;1: 235–241. doi:10.1038/681

43. Bressler DW, Silver MA. Spatial attention improves reliability of fMRI retinotopic mapping signals in occipital and parietal cortex. Neuroimage. Elsevier Inc.; 2010;53: 526–33. doi:10.1016/j.neuroimage.2010.06.063

44. Bressler DW, Fortenbaugh FC, Robertson LC, Silver MA. Visual spatial attention enhances the amplitude of positive and negative fMRI responses to visual stimulation in an eccentricity-dependent manner. Vision Res. Elsevier Ltd; 2013;85: 104–112. doi:10.1016/j.visres.2013.03.009

45. Serences JT, Yantis S. Selective visual attention and perceptual coherence. Trends Cogn Sci. 2006;10. doi:10.1016/j.tics.2005.11.008

46. Sprague TC, Saproo S, Serences JT. Visual attention mitigates information loss in small- and large-scale neural codes. Trends Cogn Sci. Elsevier Ltd; 2015; 1–12. doi:10.1016/j.tics.2015.02.005

47. Parkes LM, Marsman JBC, Oxley DC, Goulermas JY, Wuerger SM. Multivoxel fMRI analysis of color tuning in human primary visual cortex. J Vis. 2009;9: 1–1. doi:10.1167/9.1.1

48. Brouwer G, Heeger D. Decoding and Reconstructing Color from Responses in Human Visual Cortex. J Neurosci. 2009;29: 13992–14003.

49. Brainard DH. The Psychophysics Toolbox. Spat Vis. 1997;10: 433–436.

50. Watson AB, Pelli DG. QUEST: a Bayesian adaptive psychometric method. Percept Psychophys. 1983;33: 113–120. doi:10.3758/BF03202828

51. Andersson JLR, Skare S, Ashburner J. How to correct susceptibility distortions in spin-echo echo-planar images: application to diffusion tensor imaging. Neuroimage. Academic Press; 2003;20: 870–888. doi:10.1016/S1053-8119(03)00336-7

52. Smith SM, Jenkinson M, Woolrich MW, Beckmann CF, Behrens TEJ, Johansen-Berg H, et al. Advances in functional and structural MR image analysis and implementation as FSL. Neuroimage. Academic Press; 2004;23: S208–S219. doi:10.1016/J.NEUROIMAGE.2004.07.051

53. Jenkinson M, Beckmann CF, Behrens TEJ, Woolrich MW, Smith SM. FSL. Neuroimage. Academic Press; 2012;62: 782–790. doi:10.1016/J.NEUROIMAGE.2011.09.015

54. Engel SA, Rumelheart DE, Wandell BA. fMRI of human visual cortex. Nature. 1994;369: 525.

55. Swisher JD, Halko MA, Merabet LB, Mcmains SA, Somers DC. Visual Topography of Human Intraparietal Sulcus. J Neurosci. 2007;27: 5326–5337. doi:10.1523/JNEUROSCI.0991-07.2007

56. Sprague TC, Ester EF, Serences JT. Restoring Latent Visual Working Memory Representations in Human Cortex Article. Neuron. Elsevier Inc.; 2016;91: 694–707. doi:10.1016/j.neuron.2016.07.006

57. Dunn OJ. Multiple Comparisons Among Means. J Am Stat Assoc. 1961;56: 52–64.

